# RecBlast: Cloud-Based Large Scale Orthology Detection

**DOI:** 10.1101/112946

**Authors:** Efrat Rapoport, Moran Neuhof

## Abstract

**Background:** The effective detection and comparison of orthologues is crucial for answering many questions in comparative genomics, phylogenetics and evolutionary biology. One of the most common methods for discovering orthologues is widely known as ‘Reciprocal Blast’. While this method is simple when comparing only two genomes, performing a large-scale comparison of Multiple Genes across Multiple Taxa becomes a labor-intensive and inefficient task. The low efficiency of this complicated process limits the scope and breadth of questions that would otherwise benefit from this powerful method.

**Findings:** Here we present RecBlast, an intuitive and easy-to-use pipeline that enables fast and easy discovery of orthologues along and across the evolutionary tree. RecBlast is capable of running heavy, large-scale and complex Reciprocal Blast comparisons across multiple genes and multiple taxa, in a completely automatic way. RecBlast is available as a cloud-based web server, which includes an easy-to-use user interface, implemented using cloud computing and an elastic and scalable server architecture. RecBlast is also available as a powerful standalone software supporting multi-processing for large datasets, and a cloud image which can be easily deployed on Amazon Web Services cloud. We also include sample results spanning 448 human genes, which illustrate the potential of RecBlast in detecting orthologues and in highlighting patterns and trends across multiple taxa.

**Conclusions:** RecBlast provides a fast, inexpensive and valuable insight into trends and phenomena across distance phyla, and provides data, visualizations and directions for downstream analysis. RecBlast's fully automatic pipeline provides a new and intuitive discovery platform for researchers from any domain in biology who are interested in evolution, comparative genomics and phylogenetics, regardless of their computational skills.

## Findings

### Introduction

Orthology plays an extremely important role in answering a myriad of biological questions, ranging all the way from phylogenetic questions to drug target discovery. Orthologue discovery, however, is a very challenging computational task. There are several methods for discovering orthologues, among them phylogenetic, pairwise alignment and clustering techniques[1]. One of the most commonly used methods for discovering orthologues is widely known as Reciprocal Blast or Reciprocal Best Hits (RBH)[2–4]. In order for a RBH to be determined, two protein products of two different species must independently find each other as their best match in the opposite genome (see Fig. 1). In its simplest form, the Reciprocal Blast algorithm works as following: The user chooses an organism, a protein product of a specific gene from the organism's genome, and a target organism. Using the BLAST local alignment tool, the user discovers if the protein from the original organism matches a protein in the target species. If the answer is positive, the user repeats the same process, but this time by running the highest scoring match from the target organism against the original organism. If the highest scoring match is the same as the original gene, we consider it to be a successful reciprocal blast– a putative orthologue has been determined. In some cases, the definition of best hits is more flexible and allows additional candidates to be considered as orthologues [see Supplementary Fig. 2, 3, 4 in Additional file 1].

**Figure 1.**
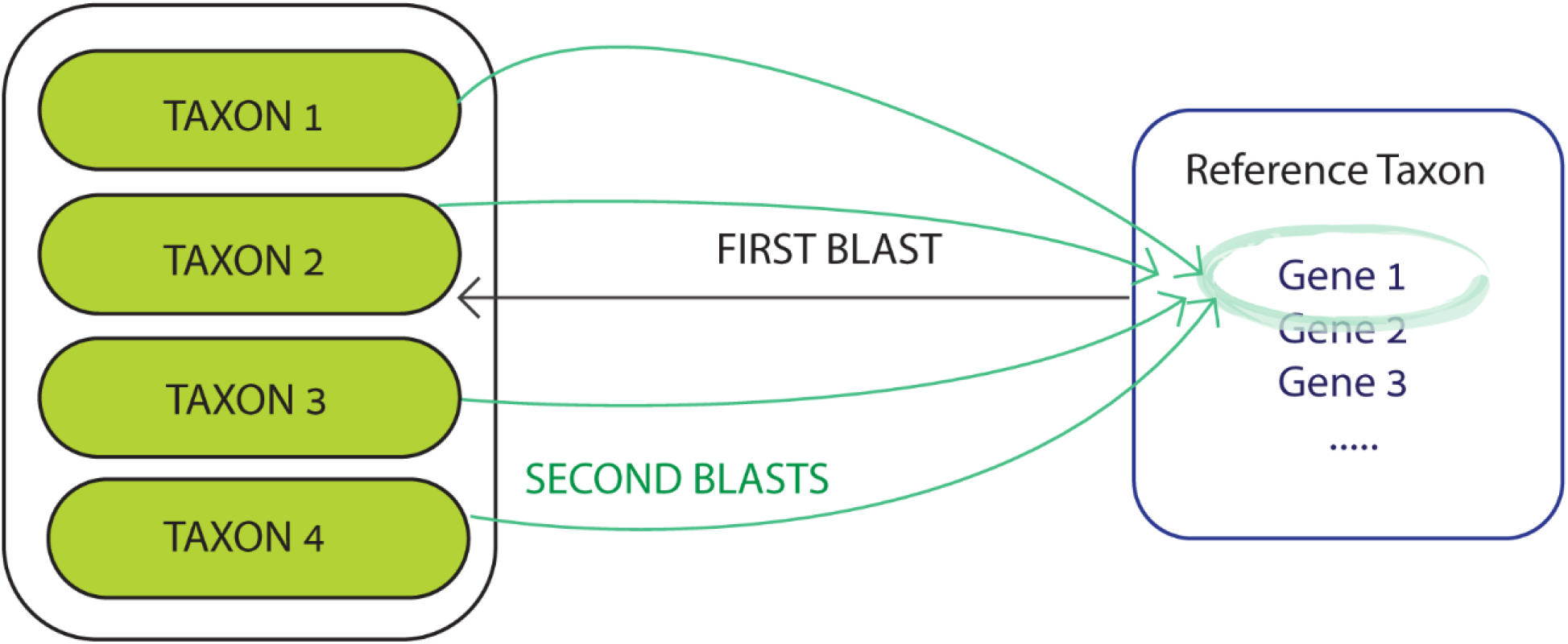
Reciprocal Blast schematic overview. Reference Taxon Gene1 is aligned to a database containing the proteins of multiple taxa. Every hit above a certain score threshold is then aligned again to the original Reference Taxon proteins.

Orthologue discovery using Reciprocal Blast is usually performed by using bi-directional BLAST[5]. This process is fairly simple when comparing a small amount of genes across two genomes. In many cases, however, researchers are not looking for orthologues between two given species, but for a broader picture spanning tens or even hundreds of taxa (in some cases even whole phyla), and hundreds of genes. A single gene alone can result in hundreds of thousands of matches to different taxa; thus, performing such a large scale comparison of Multiple Genes across Multiple Taxa (MGMT) becomes a very inefficient task. It requires complex processing of multi-dimensional comparisons (multiple taxa and genes), usually orchestrated manually. The low efficiency of this complicated, confusing and prone-to-errors process limits the scope and breadth of questions that would otherwise benefit from this powerful method.

Here we present a powerful and fully automatic pipeline, in the form of an easy-to-use cloud-based web server and a stand-alone version. Both versions are capable of dealing with complex Reciprocal Blast comparisons at scale. The interface requires as little as gene names, the taxon of origin and target taxa names in order to automatically perform and process the MGMT Reciprocal Blast in several degrees of flexibility.

Many computational tools have been developed for the purpose of orthologue detection. Each tool was designed to cater for different needs. While some allow users to investigate specific organisms from a predefined list of taxa, others allow only for specific genes to be queried, or provide specific degrees of flexibility in the orthologue discovery process[6–12]. Many of the existing tools limit the queries to dual comparisons (reference taxon against another taxon). RecBlast is a powerful, user-friendly tool that provides researchers with the ability to discover putative orthologues between any number of taxa, across any given list of genes at once, without demanding any computational background.

### Features and methods

RecBlast can be used through a flexible cloud-based web server, or by downloading a standalone version. RecBlast receives a list of parameters as input: reference taxa, a list of target organism names or taxa IDs (species of interest), a list of protein sequences the user would like to compare, BLAST parameters (e-value, identity and coverage), and a string similarity value [see Additional file 1]. First, RecBlast retrieves the query sequences, based on the user's input, and creates the respective databases. Next, RecBlast runs protein BLAST+[13] for every inspected gene, against each one of the organisms the user provided. Each result that passed the quality threshold defined by the user is blasted back against the reference organism. If the result is identical to the original gene that was used for the first BLAST, it is treated as a candidate for orthology. A result is determined identical if a) its protein sequence is similar in more than 99% to the canonical protein sequence of the inspected gene, or b) the similarity of the annotations of both proteins is greater than the 'string similarity' parameter defined by the user. The textual annotation comparison increases the chances of two proteins to be successfully matched. RecBlast works in three degrees of flexibility:

1. RBH is determined only if the very first result of the first BLAST matches the original gene when blasted back against the reference genome.
2. A “strict” match is determined for any result that passed the quality threshold in the first BLAST, if the *first* result it yields when blasted back against the reference genome is identical to the original gene.
3. The “non-strict” flexibility level counts any putative orthologue that was successfully matched to the original gene, even if it was not the first result that was yielded.

Results are provided in three separate downloadable CSV files (one for each flexibility level – strict, non-strict and RBH), each containing the number of putative orthologues found for every gene-taxon combination [see Additional file 1]. Additionally, FASTA files containing all the matches found for each sequence are provided.

### Web server implementation

RecBlast's web server is hosted and implemented on Amazon Web Services (AWS), one of the most popular cloud services available today. The server's architecture is designed to provide utmost flexibility and efficiency; the service is elastic and scales according to demand, providing an economical solution that is also stable, cost-effective and reliable. The web server is simple and easy-to-use. It requires no preparation of data, and supports various input types. The input is validated before the data is sent for processing, and the user is immediately notified on errors. The web server can receive as many as 10 different organisms and 10 different genes simultaneously per run, while the stand-alone version is only limited by the genes and taxa archived in Entrez[14], UniProt[15] and TaxDB[16] databases. Once the analysis is complete, the users receive an email with a link for their results. The web server is accessible from multiple platforms including full mobile compatibility.

### Standalone implementation

RecBlast's standalone version is open-source, and is freely available on GitHub. It can run on any machine supporting Linux with python 2.X, and supports multiprocessing. Community contributions and feedback are welcome.

### Cloud image (AMI) implementation

RecBlast's standalone version is also installed and bundled as an Amazon Machine Image (AMI) on the cloud platform of Amazon Web Services. The AMI is freely available on the AWS Marketplace and can be deployed for single use in accordance with AWS User Agreement.

### Demonstration

In order to demonstrate the potential of RecBlast, we selected 448 genes from the eukaryotic protein database (KOGs). KOGs (Eukaryotic Clusters of Orthologous Groups) are a phylogenetic classification of proteins encoded in complete genomes[17]. We used RecBlast to run the selected 448 human genes against 11 taxa, ranging from Vertebrates (*Danio rerio*) to Placozoa (*Trichoplax adhaerens*) and Porifera (*Amphimedon queenslandica*). The dataset and full analysis results, including all visualization and table-data for all three strictness levels (RBH, strict and non-strict), are provided in our Supplementary Information [see Additional file 1] and on the RecBlast website.

The results of this demonstration run show how RecBlast processes a large number of genes, across diverse and distant taxa. They demonstrate how RecBlast can assist in identifying patterns between and within groups of organisms and highlight abundant and rare or missing genes (Fig. 2 and Supplementary Fig. 8, 9 [see Additional file 1]). While results in strict mode correlate with the number of protein entries for the species in the NCBI Entrez database, RBH results show considerably weaker correlation (Supplementary Fig. 10, 11 [see Additional file 1]). The results also demonstrate how evolutionary distance influences the number of detected orthologues. Taxa distant from the inspected taxon, in this case *Homo sapiens*, are likely to have less detected orthologues. Distant taxa (*Trichoplax adhaerens, Hydra Vulgaris*), or taxa characterized by higher-frequency gene loss (*Toxocara canis*), are shown to be more distant from *H. sapiens* in terms of orthologues found. The results generated by RecBlast (visualizations, table data and FASTA sequences) can be used for further analysis of sub-groups, genes and organisms.

**Figure 2.**
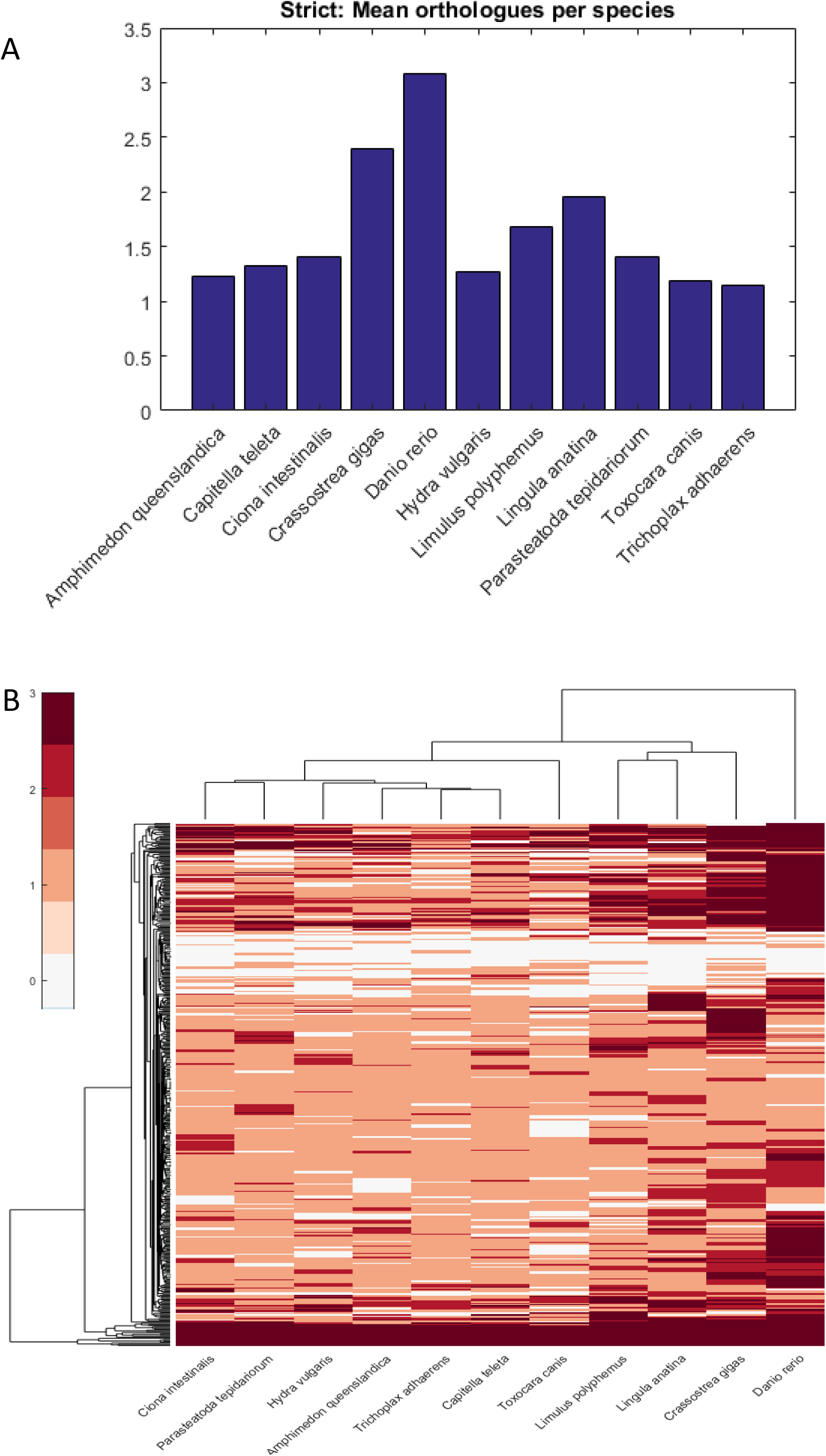
Orthologues found for the KOG genes in “strict” mode. A) The average number of orthologues found for each species. B) Hierarchical clustering of the different taxa according to the patterns of orthologues discovered.

### Discussion

RecBlast provides a fast, inexpensive and valuable insight into trends and phenomena across distance phyla, and provides data and directions for downstream analysis. RecBlast's fully automatic pipeline provides a new and intuitive discovery platform for researchers from any domain in biology who are interested in evolution, comparative genomics and phylogenetics, regardless of their computational skills.

## Availability and requirements

Project name: RecBlast

Project home page: http://www.reciprocalblast.com

Operating system(s): Web server: Any, Stand-alone version: Linux

Programming languages: Python (version 2.7)

Other requirements: Python 2.X (not 3), BLAST+ v2.3 or higher, NCBI nr database

License: MIT License

## Availability of supporting data

The data set supporting the results of this article is available in the RecBlast website (http://reciprocalblast.com/sample).

## Additional files

Additional file 1

File format: pdf

Title: Supplementary Information

Description: Background and introduction to the reciprocal Blast method, including details on the implementation of the method, and its benefits.

## Declarations

### List of abbreviations

MGMT: Multiple Genes across Multiple Taxa
RBH: Reciprocal Best Hit

## Authors' contributions

E.R. and M.N. conceived, designed and developed the system. All authors wrote and reviewed the manuscript.

## Acknowledgements

The authors would like to warmly thank Eva Jablonka for the moral support and the wise words. The authors thank Shlomi Atar for helping with our web server and the design, Barak Goldstein for his valuable support, and Oded Rechavi, Uri Ashery, Dorothée Huchon, Haim Ashkenazy and Hila Gingold for their feedback and help.

## References

1. Koonin E V. Orthologs, Paralogs, and Evolutionary Genomics. Annu. Rev. Genet. Annual Reviews; 2005;39:309–38.

2. Lake JA, Rivera MC. Deriving the Genomic Tree of Life in the Presence of Horizontal Gene Transfer: Conditioned Reconstruction. Mol. Biol. Evol. Oxford University Press; 2004;21:681–90.

3. Wall DP, Fraser HB, Hirsh AE. Detecting putative orthologs. Bioinformatics. Oxford Univ Press; 2003;19:1710–1.

4. Moreno-Hagelsieb G, Latimer K. Choosing BLAST options for better detection of orthologs as reciprocal best hits. Bioinformatics. Oxford University Press; 2008;24:319–24.

5. Altschul S, Madden TL, Schäffer AA, Zhang J, Zhang Z, Miller W, et al. Gapped BLAST and PSI-BLAST: a new generation of protein database search programs. Nucleic Acids Res. Oxford University Press; 1997;25:3389–402.

6. Sonnhammer ELL, Ostlund G. InParanoid 8: orthology analysis between 273 proteomes, mostly eukaryotic. Nucleic Acids Res. Oxford University Press; 2015;43:D234–9.

7. Aubry S, Kelly S, Kümpers BMC, Smith-Unna RD, Hibberd JM, Christin P, et al. Deep Evolutionary Comparison of Gene Expression Identifies Parallel Recruitment of Trans-Factors in Two Independent Origins of C4 Photosynthesis. Bomblies K, editor. PLoS Genet. Public Library of Science; 2014;10:e1004365.

8. Shi G, Peng M-C, Jiang T. MultiMSOAR 2.0: An Accurate Tool to Identify Ortholog Groups among Multiple Genomes. Aerts S, editor. PLoS One. Public Library of Science; 2011;6:e20892.

9. Sadreyev IR, Ji F, Cohen E, Ruvkun G, Tabach Y. PhyloGene server for identification and visualization of co-evolving proteins using normalized phylogenetic profiles. Nucleic Acids Res. Oxford University Press; 2015;43:W154–9.

10. Smith SA, Beaulieu JM, Donoghue MJ, Bininda-Emonds O, Cardillo M, Jones K, et al. Mega-phylogeny approach for comparative biology: an alternative to supertree and supermatrix approaches. BMC Evol. Biol.. BioMed Central; 2009;9:37.

11. Chen F, Mackey AJ, Stoeckert CJ, Roos DS. OrthoMCL-DB: querying a comprehensive multi-species collection of ortholog groups. Nucleic Acids Res.. Oxford University Press; 2006;34:D363–8.

12. Orsini M, Carcangiu S, Cuccuru G, Uva P, Tramontano A, Moreno-Hagelsieb G, et al. The PARIGA Server for Real Time Filtering and Analysis of Reciprocal BLAST Results. Haslam NJ, editor. PLoS One. Public Library of Science; 2013;8:e62224.

13. Camacho C, Coulouris G, Avagyan V, Ma N, Papadopoulos J, Bealer K, et al. BLAST+: architecture and applications. BMC Bioinformatics. BioMed Central; 2009;10:421.

14. Maglott D, Ostell J, Pruitt KD, Tatusova T. Entrez Gene: gene-centered information at NCBI. Nucleic Acids Res. Oxford University Press; 2004;33:D54–8.

15. Consortium TU. UniProt: a hub for protein information. Nucleic Acids Res. Oxford University Press; 2015;43:D204–12.

16. Sayers EW, Barrett T, Benson DA, Bryant SH, Canese K, Chetvernin V, et al. Database resources of the National Center for Biotechnology Information. Nucleic Acids Res. Oxford University Press; 2009;37:D5–15.

17. Koonin E V, Fedorova ND, Jackson JD, Jacobs AR, Krylov DM, Makarova KS, et al. A comprehensive evolutionary classification of proteins encoded in complete eukaryotic genomes. Genome Biol. 2004;5:R7.

